# Beyond 40 fluorescent probes for deep phenotyping of blood mononuclear cells, using spectral technology

**DOI:** 10.1101/2023.07.27.550830

**Authors:** Sandrine Schmutz, Pierre-Henri Commere, Nicolas Montcuquet, Ana Cumano, Cédric Ait-Mansour, Sophie Novault, Milena Hasan

## Abstract

The analytical capability of flow cytometry is crucial for differentiating the growing number of cell subsets found in human blood. This is important for accurate immunophenotyping of patients with few cells and a large number of parameters to monitor.

Here, we present a 43-parameter panel to analyze peripheral blood mononuclear cells from healthy individuals using 41 fluorescence-labelled monoclonal antibodies, an autofluorescent channel, and a viability dye. We demonstrate minimal population distortions that lead to optimized population identification and reproducible results. We have applied an advanced approach in panel design, in selection of sample acquisition parameters and in data analysis. Appropriate autofluorescence identification and integration in the unmixing matrix, allowed for resolution of unspecific signals and increased dimensionality. Addition of one laser without assigned fluorochrome resulted in decreased fluorescence spill over and improved discrimination of cell subsets. It also increased staining index when autofluorescence was integrated in the matrix. We conclude that spectral flow cytometry is highly valuable tool for high-end immunophenotyping, and that fine-tuning of major experimental steps is key for taking advantage of its full capacity.

## INTRODUCTION

Flow cytometry (FCM) is the most widely available method to evaluate phenotypic and functional properties (cytokine production, cytotoxicity, cell cycle) of cell suspensions (Roederer, 2002). Presently, more than 20 fluorochromes can be detected in conventional FCM although there is a foreseeable limitation in the number of fluorescent dyes that can be combined. An increasing variety of these molecules are available, many of which share similar peaks of emission, which in turn results in impossible or very high compensations. This can lead to distortions in the shape of the populations, with a risk of false positive or false negative subset identification. To obviate this problem, mass cytometry associates mass spectrometry with antibodies labelled with heavy-metal isotopes (Spitzer and Nolan, 2016). This enables combining more than 40 different parameters in a single panel. However, the cells cannot be recovered by cell sorting, the acquisition time is long and the number of events that can be analyzed is 10 to 50-fold lower than that in conventional FCM.

Spectral FCM was proposed in the 2000s and links the detection of continuous wavelength spectra with algorithms that allow the deconvolution of the data and the conversion to dot plots similar to those obtained with conventional FCM (unmixing) (Robinson, 2019). By measuring the continuous emission spectra of fluorochromes, the association of fluorescent dyes with similar peaks but different shapes of emission spectra became possible. With the first available spectral cytometer (Sony SP6800) equipped with two excitation lasers (488nm and 405nm) we built a panel of 19 fluorescent probes that allowed us to discriminate mouse splenic hematopoietic cells (Schmutz et al., 2016). More recent spectral cytometers (Cytek Aurora, Sony ID7000) are equipped with larger number of lasers and significantly increase the multiplexing capacity within a single staining panel, as has been demonstrated by a growing number of publications (Sahir et al., 2020; Park et al., 2020; Jaimes et al., 2022; Peixoto et al., 2022). Spectral cytometry provides two main advantages, the first one is the multidimensionality with high number of parameters and separation of dyes with close emission peaks (Schmutz et al., 2016), the second one is the management of the autofluorescence (AF). In a recent study we showed that incorporating AF as an independent parameter may solve difficulties in cell phenotyping and allowed the identification of new murine fetal liver cells (Peixoto et al, 2022).

In this study, we combine both features of spectral cytometry by pushing the limits of the dimensionality with the management of AF. We assembled a 43-fluorescence panel (41 antibodies, one AF parameter and one viability dye) for the analysis of human peripheral blood mononuclear cells (PBMCs). We used a new fluorochrome combination that comprised commercially available reagents and allowed the identification of most hematopoietic subsets present in PBMCs, so far described. Data analysis was greatly improved by the addition of a 320nm laser with 35 detectors, which also raised the staining index (SI) of some populations. We distinguished 28 sub-populations in an unsupervised analysis using a dimensional reduction FIt-SNE that matched the primary subsets mentioned in the PBMCs. We conclude that spectral FCM enables the examination of many factors and has the potential to increase analytical capability as new fluorescent probes are continuously developed.

## MATERIALS AND METHODS

### 1. Preparation of cells

Human Blood (50 mL) from healthy donors was collected on heparin tubes by ICAReB facility of the Institut Pasteur, Paris. Peripheral blood mononuclear cells (PBMCs) were isolated by density gradient centrifugation (Ficoll-Paque™ Premium, Dutscher, Brumath, France), washed twice, counted and resuspended in Cell Staining Buffer (BioLegend). Cells were counted using a Countess II™ (Thermo Fisher Scientific) and 3 million cells were taken for staining with the cocktail of antibodies.

### 2. Cell staining and preparation of controls

All antibodies (Table 1) were titrated to give the highest signal-to-noise ratio. The panel was first tested with 38 antibodies (all antibodies except the four custom dyes). PBMCs were stained in four steps (Table 2) of 20 minutes each, all at 4°C, with one washing step with Cell Staining buffer (BioLegend) between each group of antibodies. The sequential staining was done in 5 mL FACS tube and after the last centrifugation, cells were resuspended in 400 µl of Cells Staining Buffer and transferred into a 96-half-deep well plate. 1 million unstained PBMCs were used as negative control for the staining and for AF detection.

**Table 1:**
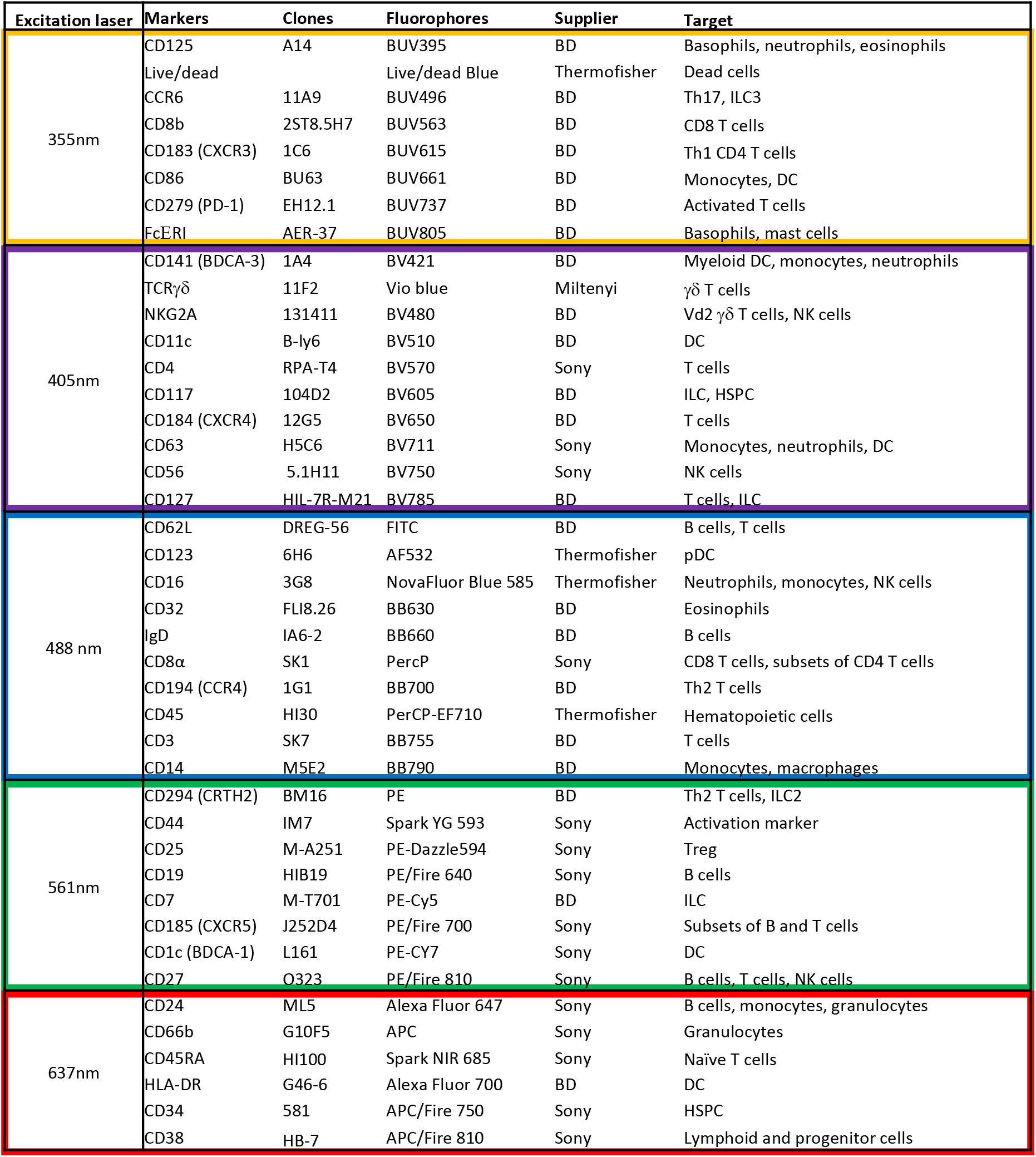
42-color panel for PBMC deep phenotyping. Antigens and clones used in the 42-color panel designed for the 6-laser spectral analyzer ID7000, in order to phenotype freshly isolated human PBMCs. Abbreviations: Th: T helper, ILC: Innate Lymphoid Cell, DC: Dendritic Cell, pDC: plasmacytoid Dendritic Cell, NK: Natural Killer, HSPC: Hematopoietic Stem and Progenitor Cell, Treg: Regulatory T cell.

**Table 2:** The staining of the PBMCs has been performed in 4 steps. The antigens of each group are listed.

| Group 1 | Group 2 | Group 3 | Group 4 |
| --- | --- | --- | --- |
| CD125 | CD62L | CXCR3 | CD8a |
| CCR6 | CD123 | CD86 | CD66b |
| CD8b | CD32 | FcERT | CD24 |
| PD1 | IgD | BDCA3 | CD45RA |
| CD63 | CCR4 | NKG2A | HLA-DR |
| CD56 | CD3 | CD11c | CD34 |
| CD127 | CD25 | CD4 | CD38 |
| TCRgd | CD19 | CD117 | CD45 |
| CXCR5 | CD7 | CXCR4 |  |
| CRTH2 | BDCA1 | CD44 |  |
| CD16 | CD27 | CD14 |  |
| Live dead |  |  |  |

Single staining controls were prepared on UltraComp eBeads Plus (ThermoFisher Scientific), incubated 20 minutes at 4°C with the conjugated antibodies, washed with Cell Staining buffer and resuspended in 400 µl of Cell Staining buffer. Viability dye single staining was prepared on ArC™ Amine Reactive Compensation Bead Kit (ThermoFisher Scientific).

### 3. Cytometer set-up and sample acquisitions

Stained cells and controls were acquired on a Sony ID7000 spectral cytometer. This cytometer is configured with 6 lasers (320nm, 355nm, 405nm, 561nm, and 637nm) and the PMT voltages were set up for each laser independently. Before acquisitions, quality control of the instrument was done using align check and 8-peak beads, following the procedure of the instrument user guide. A maximum of cells was recorded for each stained sample, between 1 and 1,6 million cells. For the single stainings, a minimum of 2000 beads were recorded.

### 4. Data analysis

Unmixing was done on the ID7000 software. For each dye of the panel, negative and positive spectrum shape were assigned before unmixing was calculated. Unstained cells were analyzed with the Autofluorescence Finder tool. Autofluorescent cells were gated and added to the unmixing. In total, 43 parameters were unmixed. Unmixing quality was verified using the Unmixing Viewer tool and small adjustments, if needed, were done using the Spectral Adjuster tool.

Unsupervised data analysis was performed using an integrated Sony solution, in collaboration with the Sony R&D team.

## RESULTS

### 1. Antibody panel construction

The number of commercially available dyes is constantly increasing with variable SI, therefore the panel design is crucial to get the better resolution between the markers. As a result, the antibody panel is the most important element for a high dimension FCM analysis. Table 1 shows the combination of dyes and antibody targets used in the panel designed for this study. We initially selected 35 fluorescence-labelled monoclonal antibodies that were titrated and tested individually on beads and on cells. Since two did not show a detectable staining when used together, the fluorophores of these two antibodies were modified in the final version of the panel. Once the 35-antibody panel was tested, we added two-by-two 7 additional fluorescence-labelled antibodies. To improve resolution and decrease background, we implemented a 4-step staining (Table 2). The less expressed antigens or dim markers were stained in the first step (group 1), followed by staining by larger dyes and/or of highly represented markers in later steps. To equilibrate the overall staining, similar number of antibodies was used in each step. Once implemented, this staining procedure was reproduced in three independent experiments performed on cells of two different donors.

Figure 1 shows the spectral emission of each fluorochrome across the individual detectors. The dyes excited by the 355nm UV laser (A) were mostly detected in the 320nm and in the 355nm detectors, the dyes excited by the 405nm violet laser (B) were also found in the 355 nm detector. The dyes excited by the 488nm blue laser (C) were found across all detectors showing the highest degree of cross excitation, whereas excitation by the 561nm yellow/green laser (D) was also found in the blue detectors. Finally, dyes excited by the 637nm red laser (E) showed the lowest degree of cross excitation.

**Figure 1:**
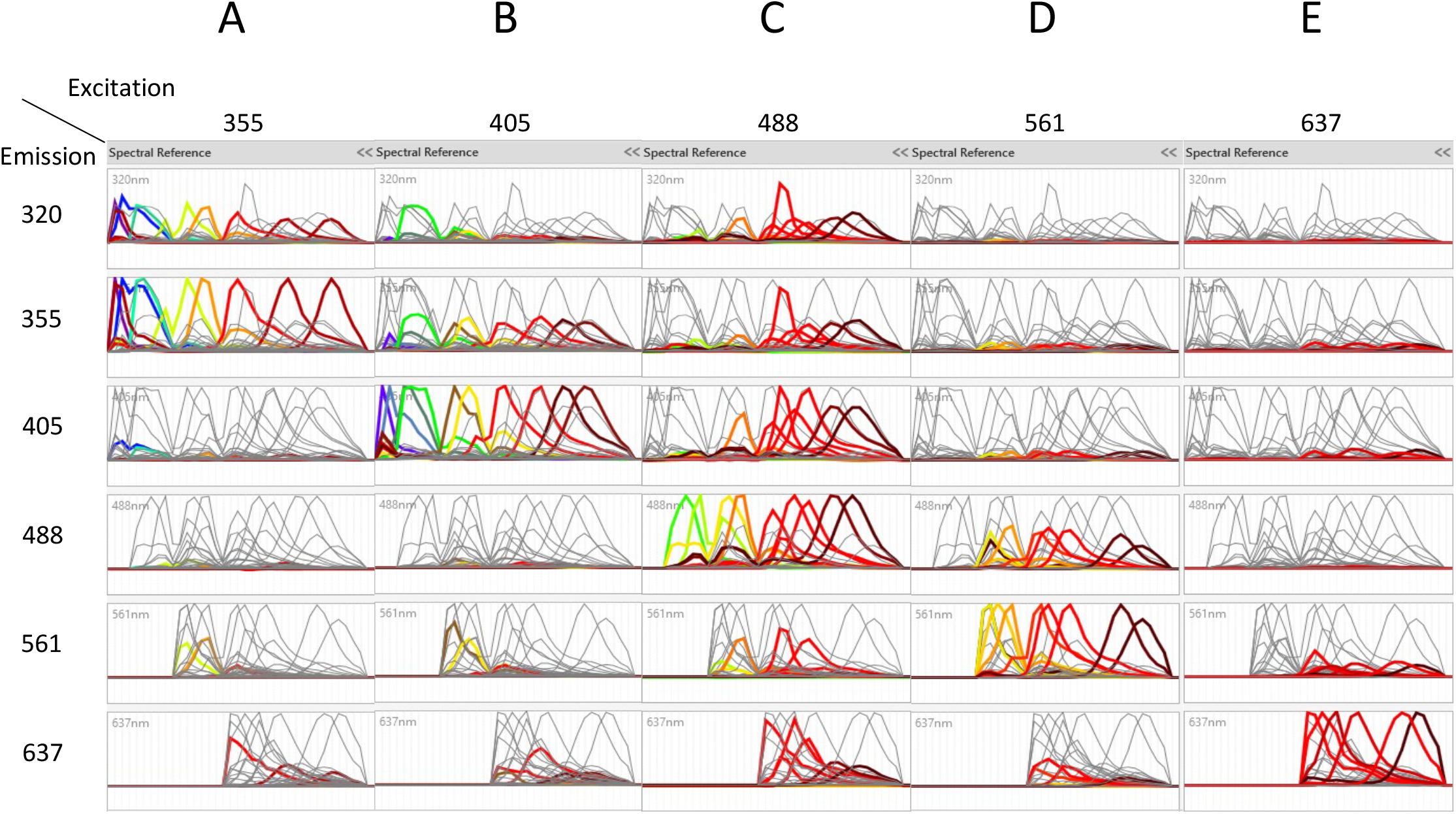
The spectra of the different dyes are shown on the detectors of all lasers. Emission spectra after unlmixing of the dyes excited by the 355nm laser (A), the dyes excited by the 405nm laser (B), the dyes excited by the 488nm laser (C), the dyes excited by the 561nm laser (D), and the dyes excited by the 637nm laser (E) are shown on the detectors of all lasers. In each column, each color represents a separated dye.

In spectral cytometry, the similarity matrix indicates close emission peaks and/or very similar spectral signatures and determines the capacity to separate dyes. Modifications of fluorochrome combinations may have a significant impact on the total similarity index of the panel. The combination of fluorochromes in the panel shown in Table 1 was such that no similarity index higher than 0.85 was allowed, and this resulted in minimal distortion (Sup Figure 1). Two by two plots in the combinations (SparkNIR-685/AF647; PE-Fire640/PE-Cy5 and BV510/BV480) show that despite the highest similarity index, populations are correctly discriminated. These results confirmed that a careful choice of fluorochromes in the construction of the antibody panel is the most important element for optimal discrimination of the populations with a complex panel.

### 2. Autofluorescence management removes phenotyping artefacts

The unstained sample of PBMCs was analyzed using the AutoFluorescence Finder available on the ID7000 software (Futamura et al., 2015). This tool allows the visualization of unspecific fluorescent signal(s), present in the unstained sample, along the lasers (Sup Figure 2). This signal is gated and presented as AF to the software that uses it as a supplementary parameter when calculating the unmixing. With PBMCs, one population of AF was found and added to the 42-color panel, thereby transforming it into a 43-color panel comprising all fluorescent dyes and the AF (Sup Figure 2). The autofluorescent population was clearly detected in the 320nm, 355nm and 405nm lasers and to a lesser extent in the remaining lasers (Figure 2A). The spectrum of AF is overlaying almost completely with the live/dead FVS440 spectrum (Figure 2A). Failing to take the AF parameter into account results in 3 populations of various intensities of live/dead marker and several distortions (Figure 2B, left plot). We verified the impact of the matrix calculated with or without the AF parameter on all the proportions of downstream populations and including the autofluorescent parameter removed the populations with different intensity of viability dye (Figure 2B, middle plot). AF identifies subpopulations of CD14^+^ myeloid cells (Figure 2B, right plot) that constitute the majority of circulating monocytes, but also neutrophils and dendritic cells.

**Figure 2:**
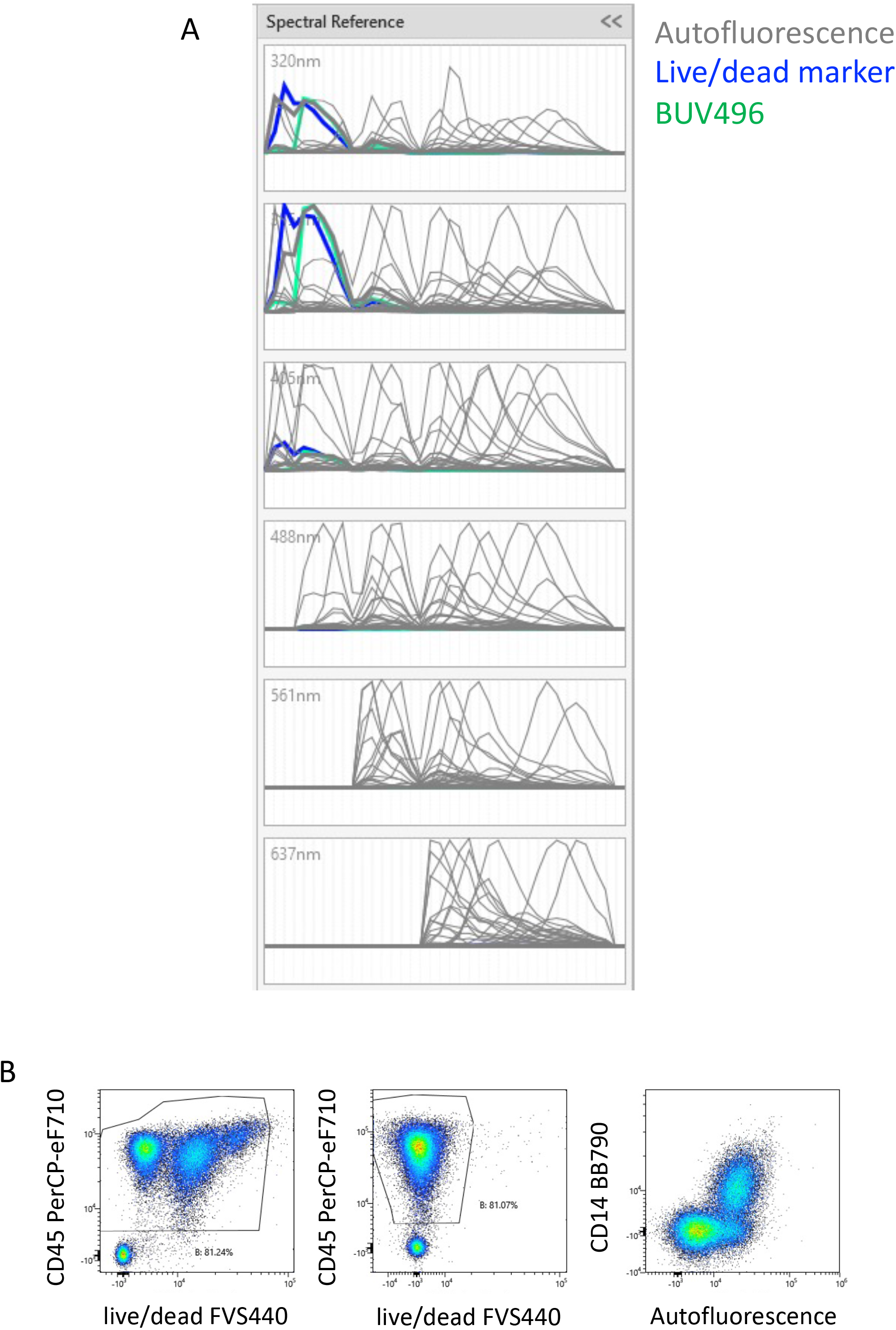
Effect of AF management on the unmixing. **A**. Analyzing unstained PBMCs with the Autofluorescence Finder tool allows the definition of an autofluorescent population with a spectrum (in grey) that highly overlays the live/dead marker (in blue). **B**. Without considering the AF, cells with three intensities of live/dead marker are visible (left plot). By including the autofluorescent population as a parameter, the artefact of the different levels of live/dead marker is absent (middle plot). Most of the autofluorescent cells are CD16^+^ monocytes (right plot).

The incorporation of the 320nm laser in the unmixing matrix has a surprising impact on the quality of the analysis because it increases the discrimination between live/dead and AF. Whereas the unmixing using only 5 lasers (355nm-637nm) shows population distortions (Figure 3A and Sup Figure 3A), the introduction of the data measurement by an additional laser (320nm) resulted in an optimal NxN matrix and consequent increased SI for several populations (Figure 3B and Sup Figure 3B). Figure 3 shows the examples of BUV563 vs BUV496, BV421 vs BUV496, and BUV395 vs BUV737 that have reduced spill over and spread upon addition of 320nm excitation and AF management (Figure 3B). As particularly relevant, we marked with green arrows three examples of low SI that were increased when the 320nm laser is switched on (6 lasers configuration) (Sup Figure 3A and 3B).

**Figure 3:**
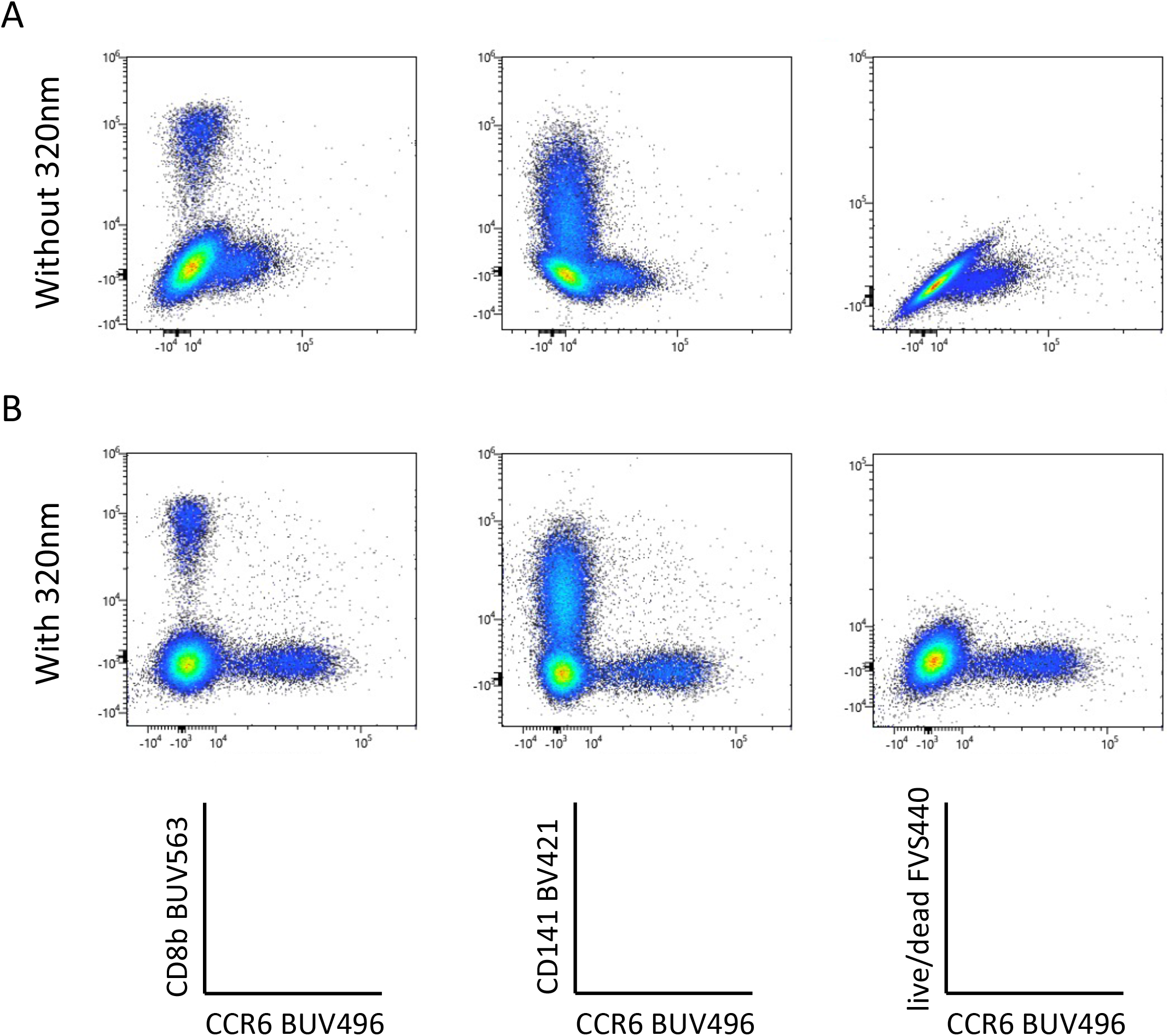
The resolution of several combination on bi-parametric plots is highly increased when the 320nm deep-UV laser is used (B) compared to analysis done without the 320nm laser (A).

To verify whether the AF parameter has an impact on the unmixing even when there is no excitation with the 320nm laser, we then compared the staining profiles whether or not AF management is included in the unmixing in the 5 lasers configuration (no 320nm laser). In the absence of the sixth laser SI are better if no AF management is introduced in the analysis (Sup Figure 3C), although artifactual populations, corresponding to AF are now detected (purple arrow, Sup Figure 3A, B, and C). Our results show that in the 5 lasers configuration, SI are higher if the AF management is not included in the analysis, but on the other hand the presence of AF expressing artificial populations was detected. Increase in the number of excitation lasers was decisive factor to improve the quality of the analysis.

### 3. Unmixing validation

Unmixing of 42 colors was done by assigning positive and negative populations in single stained samples (Sup Figure 4A) and verified in two steps, first by analyzing the shapes of different dyes (Figure 1). Interestingly, almost only the dyes excited by the 637nm laser were generally not co-excited by other lasers, as shown by the flat signal on the other lasers. In contrast, most of the dyes excited by the 355, 405, 488 and 561nm lasers were co-excited by other lasers, which helped in separation and better resolution of close dyes.

In a second step, validation of unmixing was done by looking at the NxN matrix showing all the combinations (Sup Figure 4B). Where required, we used the spectral reference adjuster tool to finely adjust the population discrimination (Futamura et al., 2015).

### 4. Data analysis

Manual gating allowed the characterization of all CD45 expressing populations of PBMCs (Park et al., 2020). Due to the complexity of the analysis, we initially subdivided cells into CD3 expressing T cells, CD19 expressing B cells and CD3^-^ CD19^-^ cells (Figure 4).

**Figure 4:**
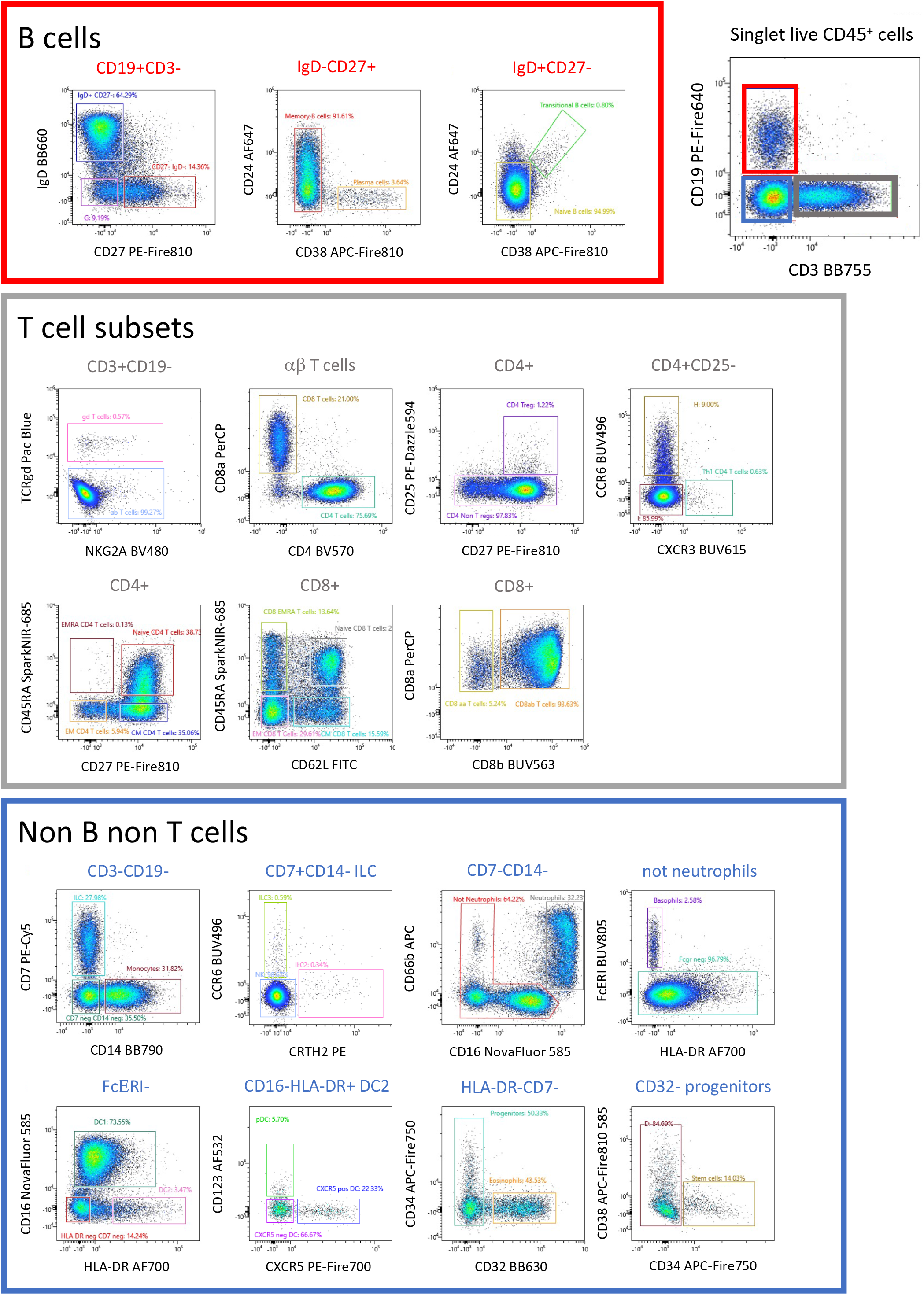
This high-dimension panel allows the analysis of most populations of lymphoid and myeloid populations present in PBMCs. This manual analysis started with the separation between B cells, T cells and non-B non-T cells.

Increasing the number of parameters analyzed on one sample increases the number of possibilities for the gating strategy and manual gating is time-consuming and biased way of analyzing data. In this study we performed in parallel manual gating and unsupervised analysis.

We used a FIt-SNE dimensional reduction tool that subdivided all major subsets in an unsupervised manner (Figure 5A), after manual gating on singlet live CD45^+^ cells. Subsequent analysis of different T cell subsets allowed the identification of αβ and γδ expressing cells, as well as CD4 and CD8 expressing T cells (Figure 4, Figure 5B). A deeper analysis of these cells identified naïve, central memory (CM), effector memory (EM) and effector memory RA (EMRA) expressing CD4 and CD8 T cells. Within the CD4 cells we identified CD25-expressing Treg (Figure 5C), CXCR3^+^ Th1, CCR6^+^ CXCR5^-^ Th17 and CXCR5^+^ follicular helper T cells (Tfh). CD27 and CD62L were interchangeable for the definition of naïve and CM populations in both CD4 and CD8 subsets. The B cell compartment could be subdivided into naïve and memory B cells, plasma cells and transitional B cells.

**Figure 5:**
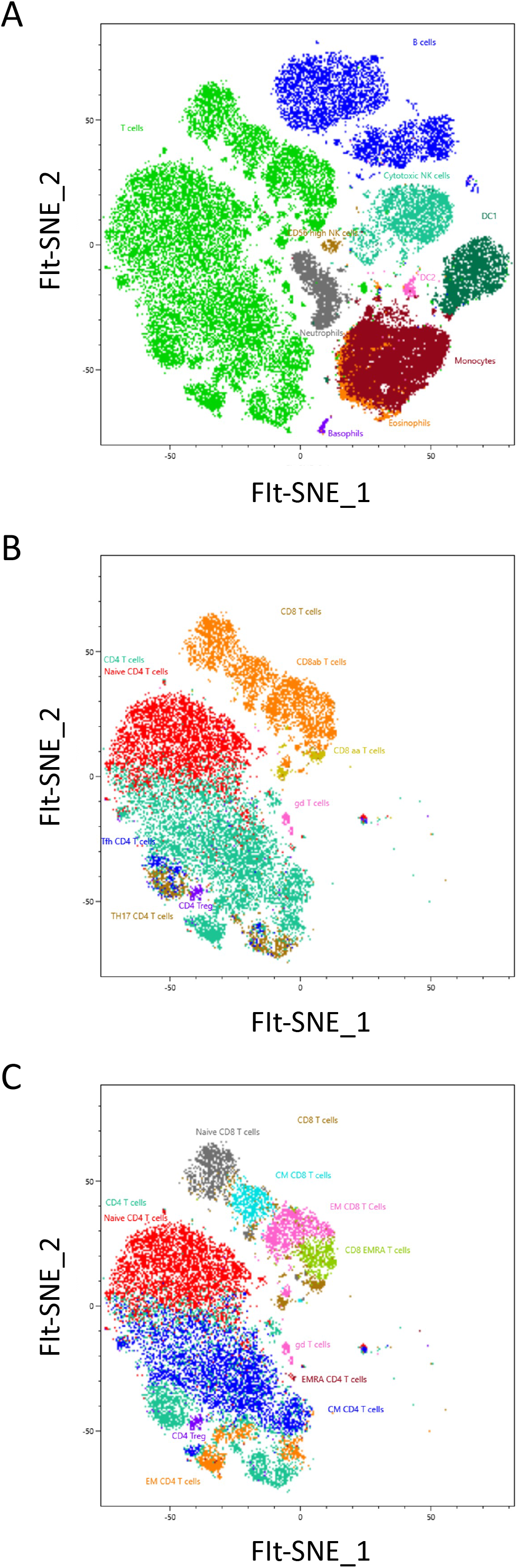
The analysis has been completed with a Sony-based tool for unsupervised analysis. The unsupervised analysis has been run from singlet live CD45^+^ cells. Shown are FIt-SNE representations, first of the main populations contained in the PBMCs (A), then of the first level of T cell subpopulations (B), and finally with all the small subsets of T cells (C).

In the non-T non-B compartment, CD14^+^ monocytes were the dominant population followed by CD7-expressing ILC that comprise NK cells (CD56-expressing and CD16-expressing killer NK), ILC3 and ILC2 (Figure 4, 5). All subsets of HLA-DR^+^ DC (CD16^+^ DC1, CD123^+^ pDC, as well as CXCR5-expressing DC) were detected. Neutrophils (CD66b, CD16^+^), eosinophils (CD32^+^) and basophils (FcγR^+^) were also clearly detected, as well as a minor population of CD34^+^ CD38^-^ stem and progenitor cells and CD38^+^ CD34^-^ progenitors. Taken together our strategy allows identifying the majority of immune cell populations in PBMCs, comprising several subsets of circulating B cells, T cells, ILC, DC, 3 populations of granulocytes and two minor populations of progenitors. The outcome was similar between both manual and unsupervised methods, with deeper phenotyping in the unsupervised manner.

## DISCUSSION

We assembled here a 41-fluorescent antibody panel that additionally include an autofluorescent parameter and a viability dye. This panel has similar complexity to previously reported high-end panels using different technologies (Spitzer and Nolan, 2016). Large antibody panels used in spectral FCM were also previously reported (Sahir et al., 2020; Park et al., 2020). These antibody panels included fluorescent dyes with similarity index higher than 0.9 that resulted in population distortions that compromised subset identification (Sahir et al., 2020). The panel assembled here indicates that keeping similarity indexes between fluorochromes below 0.85 improves the resolution of the populations and avoids spill over. The detection of AF was essential even when analyzing populations of circulating hematopoietic cells, where AF is not usually a concern. Two aspects can be highlighted: 1. AF integration in the unmixing deduces the AF from the real antibody staining thus preventing artifacts 2. It offers a new parameter that increases the analytical power. In tissues with high levels of AF across multiple cell populations, we have previously demonstrated that this parameter simply allows the discrimination of specific populations, demonstrating the analytical capacity of spectral analysis (Peixoto et al., 2022). In the present study we analyze a population of blood PBMCs, largely depleted of neutrophils. We could identify AF in a subset of myeloid cells that we used after integration in the unmixing matrix as an additional analytical parameter. This allowed the identification of most subsets of B and T lymphocytes that are well represented in PBMCs (15% and 47%, respectively). We could also detect different subsets of NK cells and rare innate lymphoid cells (ILC2). In addition to monocytes, that are an abundant fraction of PBMCs (30% of CD45^+^ non-B non-T cells), we detected two subsets of dendritic cells, plasmacytoid and myeloid, as well as basophils and eosinophils, that comprise less than 0.1% of the circulating cells. Neutrophils are the most abundant myeloid subset in circulation accounting for 40% to 70% of the PBMCs in whole blood, but their frequency is severely reduced by the isolation procedure that include a density gradient. In these samples we detected around 3% of neutrophils, identified as CD3^-^ CD19^-^ CD66b^+^ CD16^+^ cells. Importantly, we detected rare populations of stem and progenitor cells as CD117^+^ CD34^+^ and CD38^+^CD34^-^ cells, respectively, that represent less than 0.01% of the circulating cells.

Interestingly, we found that an additional laser in the deep UV spectrum, 320nm laser, that did not have an assigned excitation function, cross excited UV (355nm)- and blue (488nm)-excited fluorochromes. Integration of this data in the unmixing resulted in a significantly better resolution of the positive signal and consequently increased the SI of several cell populations allowing for a better discrimination of rare subsets, including stem and progenitor cells. Interestingly, in the absence of the 320nm laser, distortions and decreased SI were avoided by excluding the AF parameter from the analysis indicating that AF management does not always improve the quality of the data. This result indicates that, for large panels, equipping analyzers with increasing number of lasers substantially improves the quality of the data. We conclude that spectral flow cytometry, with its capacity to unmix dyes with very similar spectral emission signatures and the fact that AF is considered in the unmixing, is the technology that provides the best analytical capabilities with user friendly handling that does not require dedicated personnel and with unlimited potential, as new fluorescent probes that become available in the market can be readily incorporated in the panels.

## FIGURE LEGENDS

**Supplementary Figure 1:** Unmixing of dyes with high similarity index. Singlet live CD45^+^ populations are shown after unmixing. **A**. AF647 and SparkNIR685 have a similarity index of 0.84 **B**. PE-Cy5 and PE-Fire810 have a similarity index of 0.77 **C**. BV480 and BV510 have a similarity index of 0.78.

**Supplementary Figure 2:** AutoFluorescence Finder tool in ID7000 software. **A**. Definition of the autofluorescent population using the Autofluorescence Finder tool of the ID7000 software. The spectral shape of the autofluorescent population is shown in blue.

**Supplementary Figure 3:** Impact of 320nm deep-UV laser. Comparison of the signal resolution and spill over upon acquisitions of the cells with 5 lasers (A, C) or 6 lasers, including 320nm laser (B) and analysis of the generated data with AF management (A and B) or without (C). The example of the expression of BUV496 signal vs each fluorophore of the panel is shown.

**Supplementary Figure 4:** Unmixing was validated with the NxN matrix showing all bi-parametric plots for one dye after the other. **A**. Example of assignment of positive and negative population from single stained sample of PerCP (left plot). The ribbon plot shows the spectrum of the positive gate of PerCP all along the detectors of all lasers. **B**. NxN matrix with all combinations is a tool to verify and validate the quality of the unmixing for each dye, one after the other, versus all the other dyes. Here is shown PerCP versus all other dyes of the panel.

