## Supplementary figures and images for "Beyond 40 fluorescent probes for deep phenotyping of blood mononuclear cells, using spectral technology"

### Sup Figure 1

#### Supplementary Figure 1

A

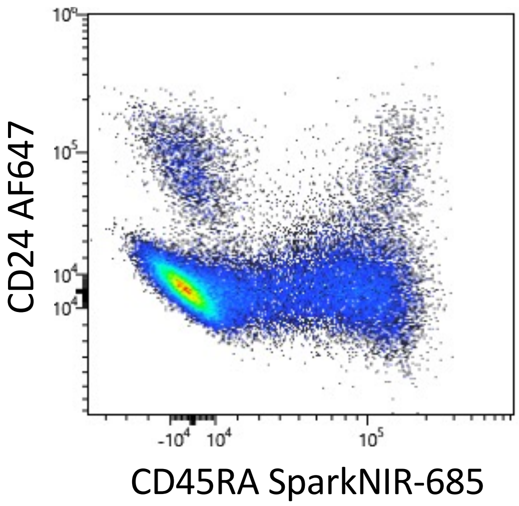


Similarity Index 0.84

B

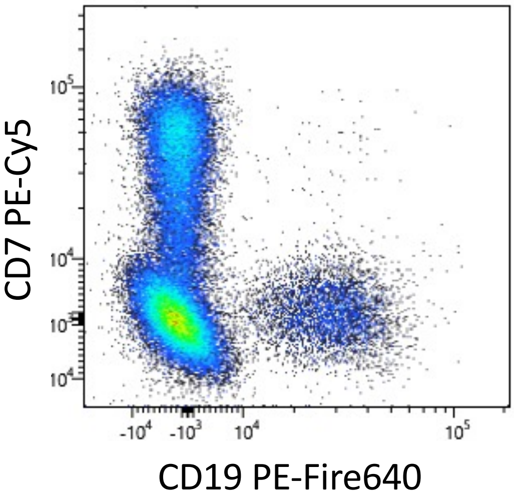


Similarity Index 0.77

C

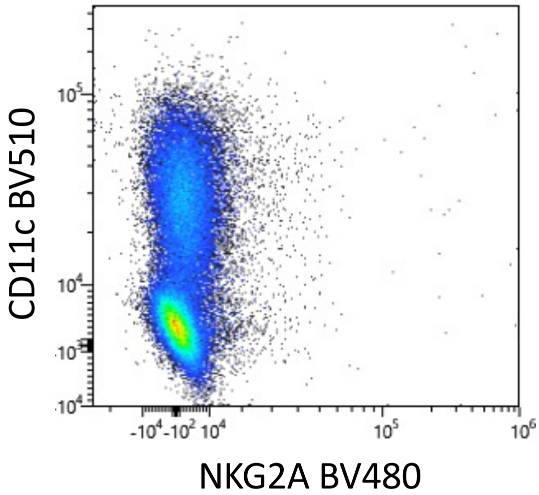


Similarity Index 0.78

### Sup Figure 2

#### Supplementary Figure 2

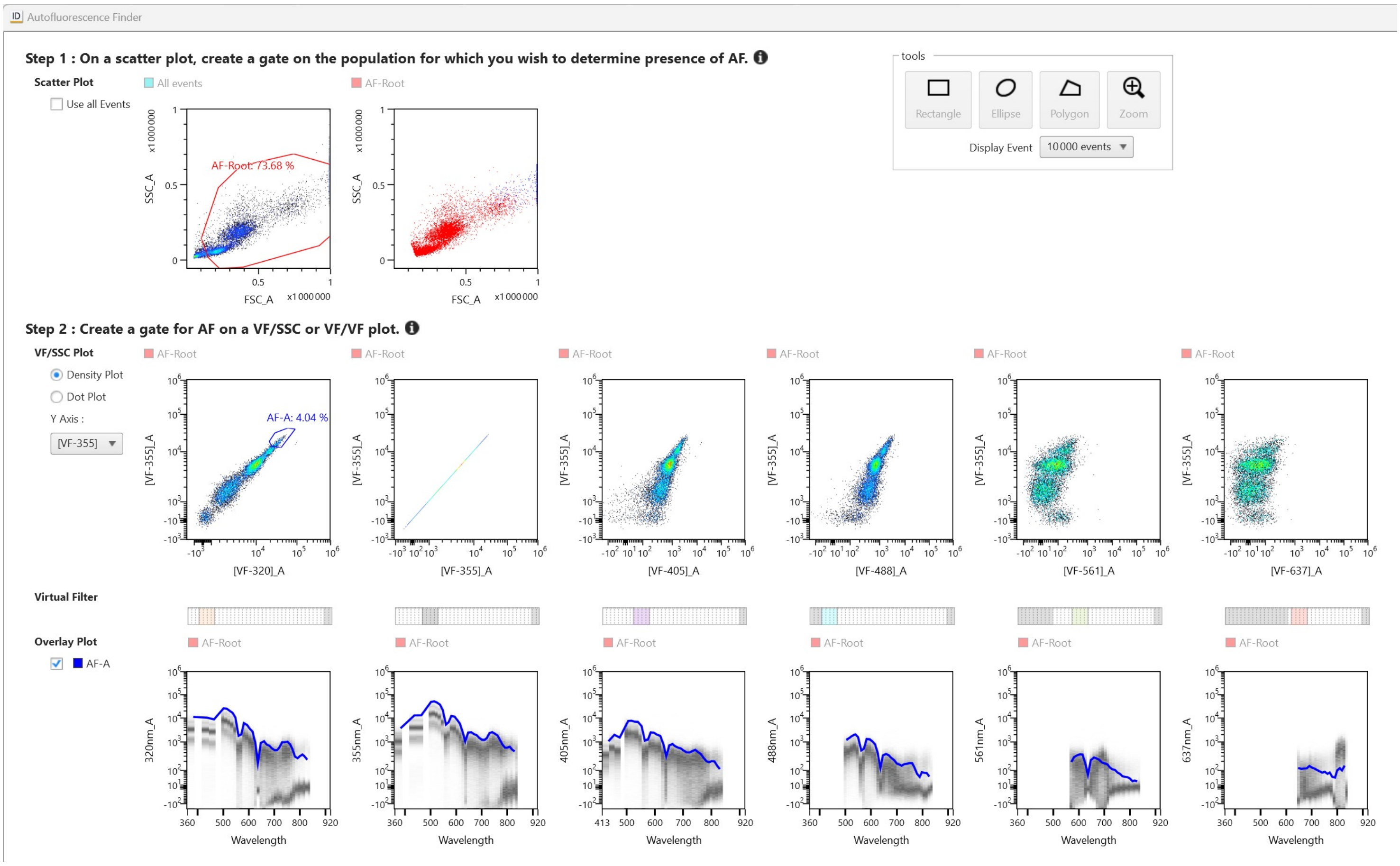

### Sup Figure 4

#### Supplementary Figure 4

A

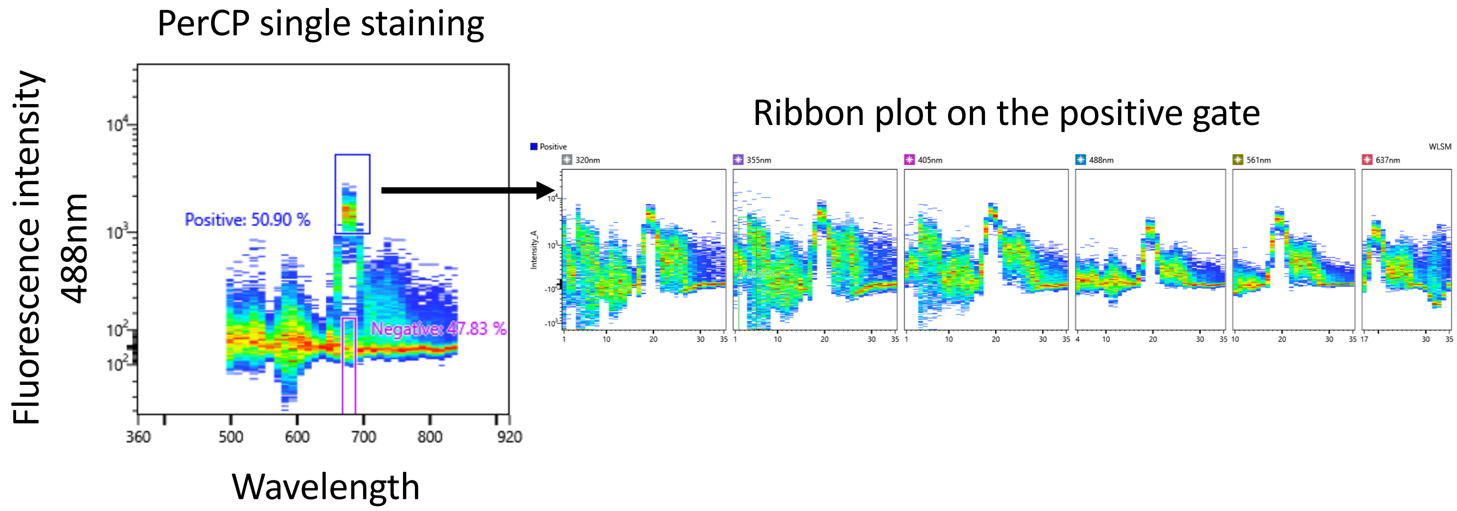


B

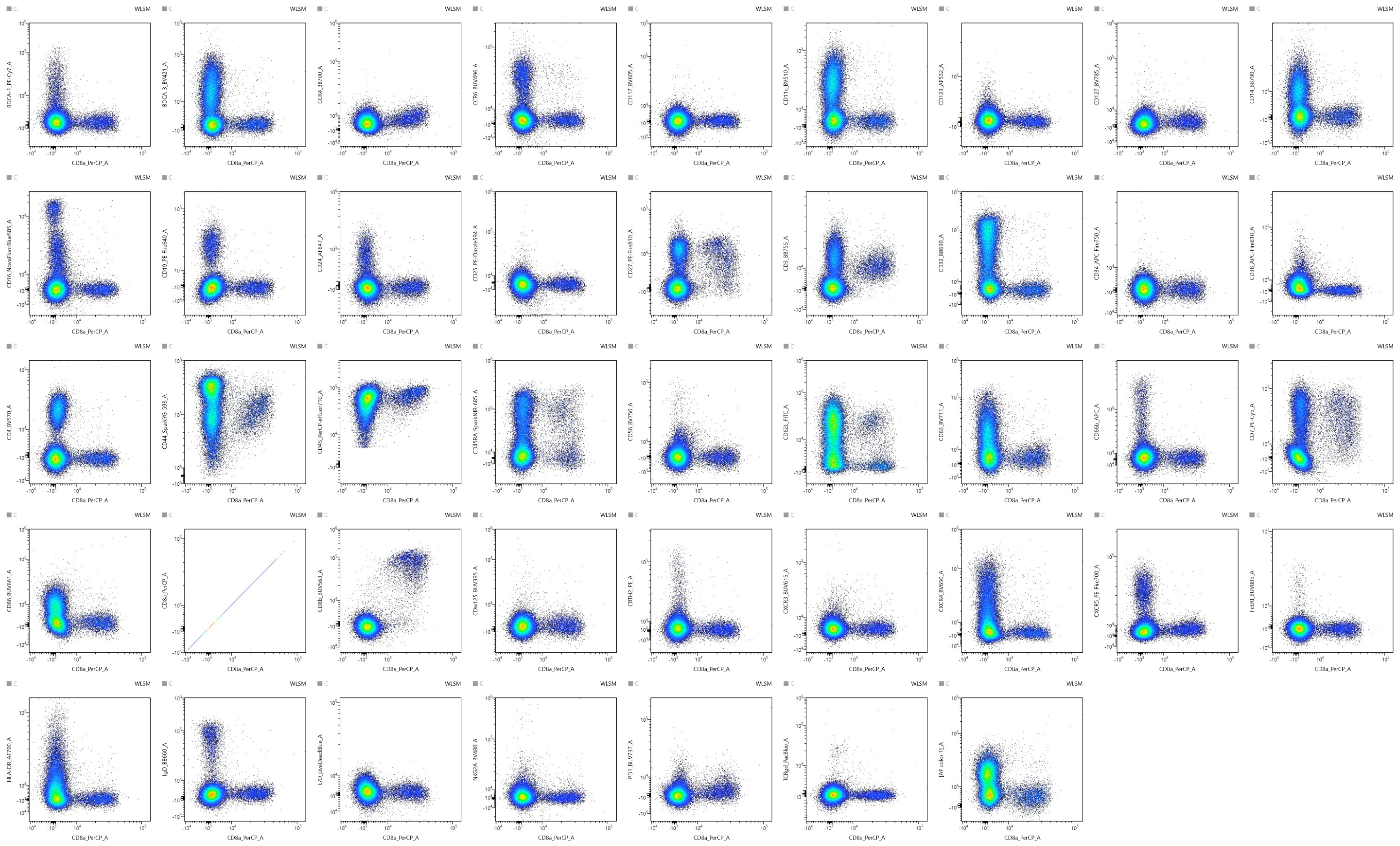
