## Supplementary material for "Beyond 40 fluorescent probes for deep phenotyping of blood mononuclear cells, using spectral technology": Sup Figure 3

### Supplementary Figure 3

#### A 5 lasers + AF management

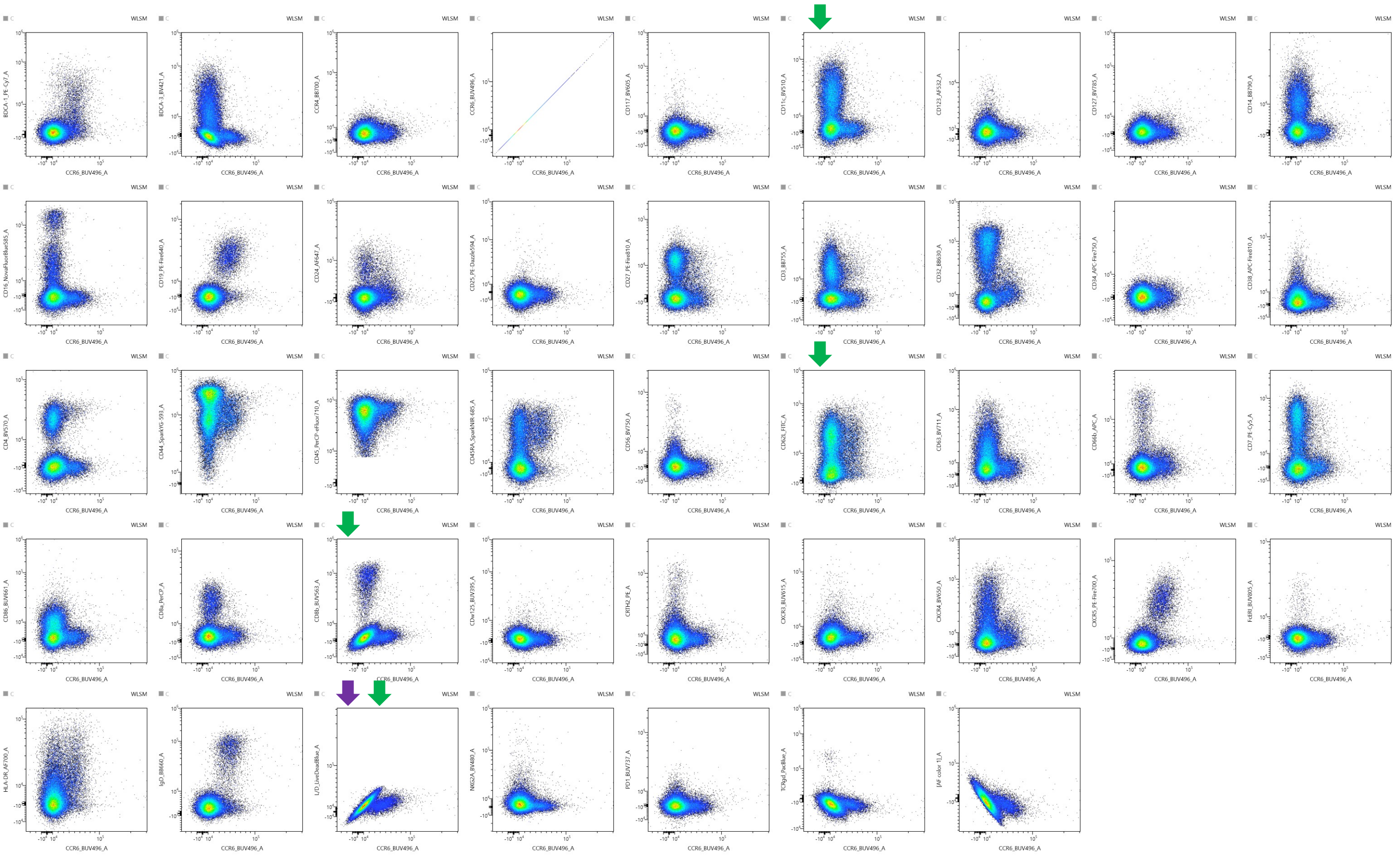

#### B 6 lasers + AF management

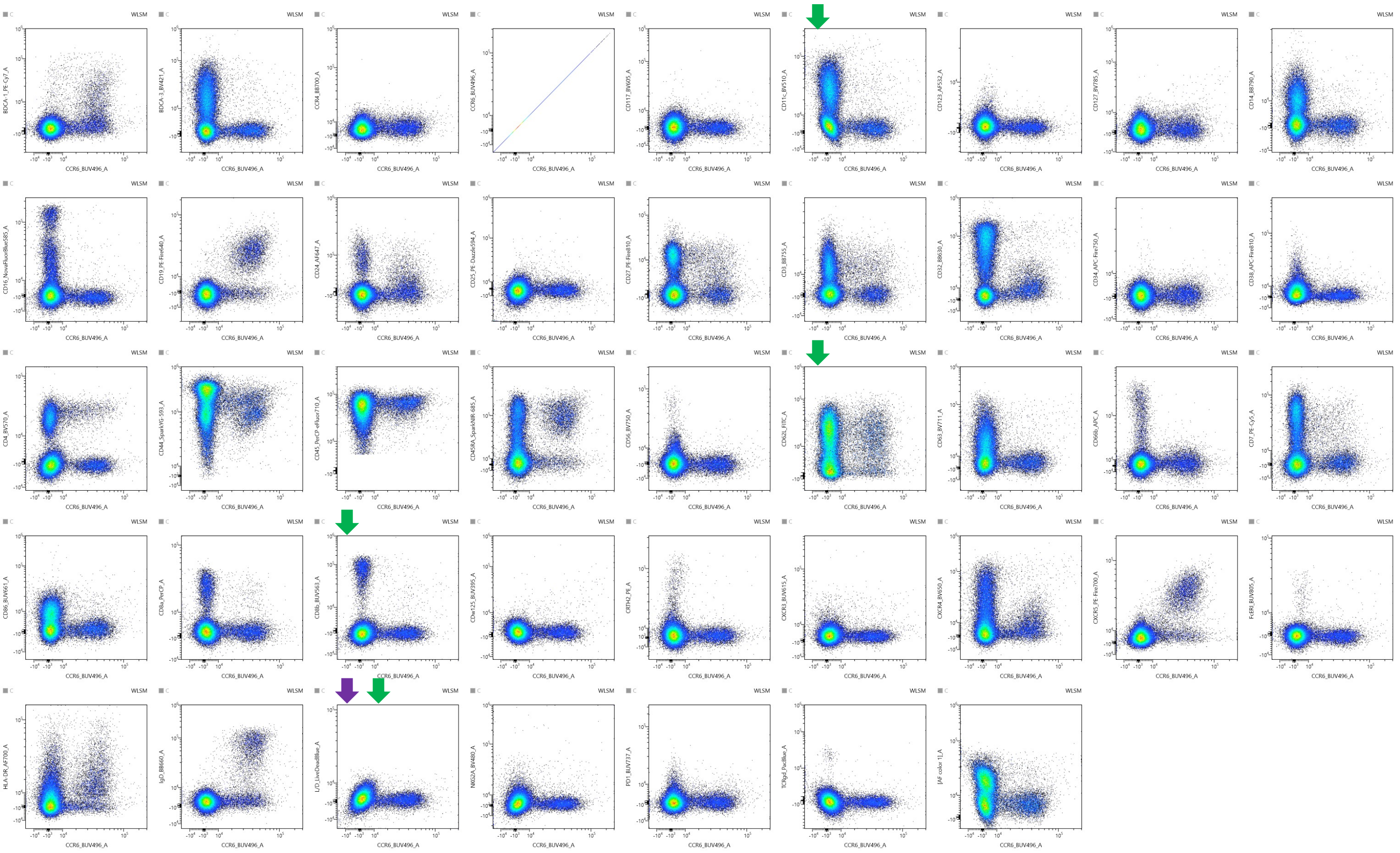

### Supplementary Figure 3

#### C 5 lasers no AF management

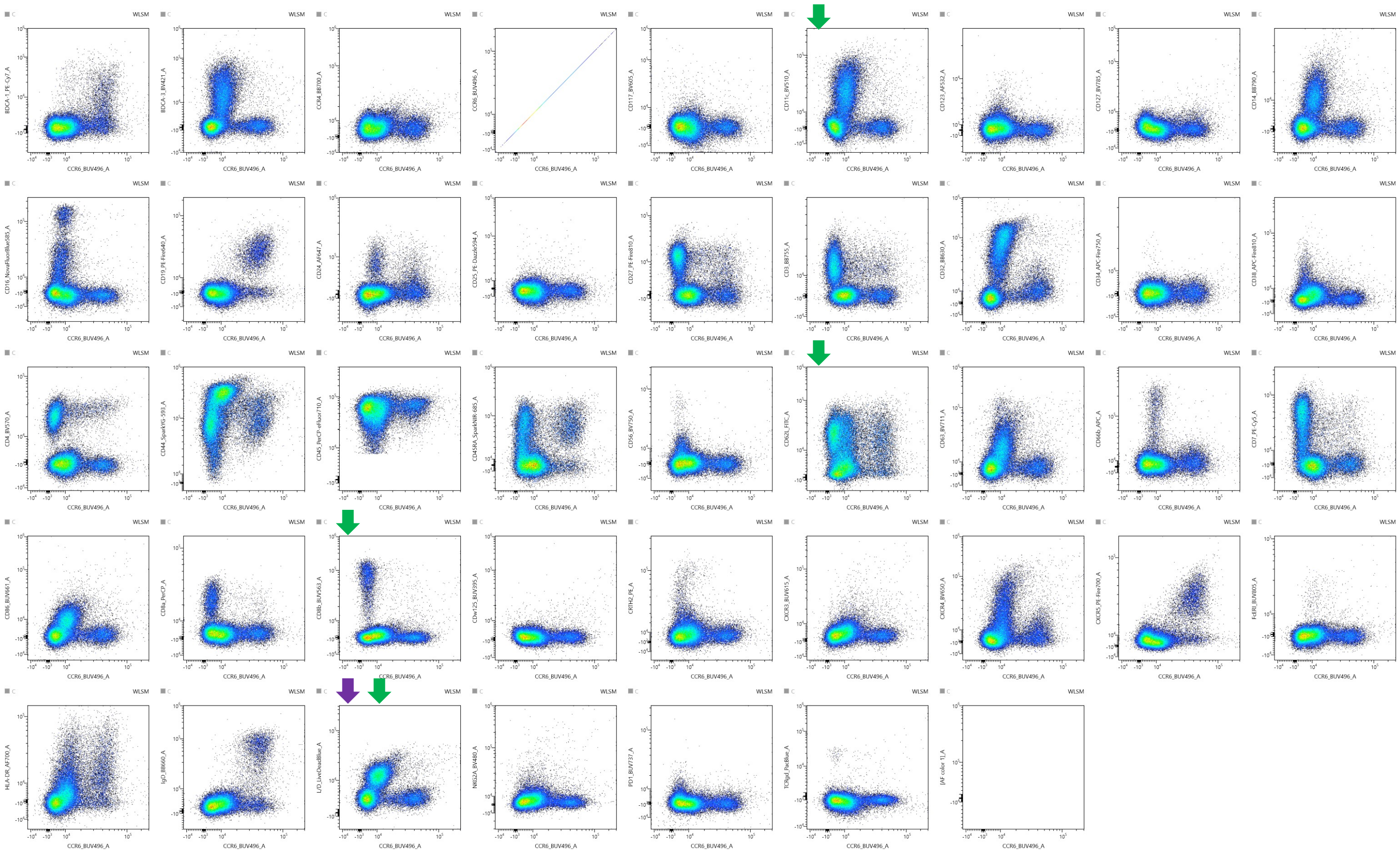
